# Enhancing CRISPR deletion via pharmacological delay of DNA-PK

**DOI:** 10.1101/2020.02.12.945907

**Authors:** Núria Bosch, Michaela Medová, Roberta Esposito, Carlos Pulido-Quetglas, Yitzhak Zimmer, Rory Johnson

## Abstract

CRISPR-Cas9 deletion (CRISPR-del) is the leading approach for eliminating DNA from mammalian cells and underpins a variety of genome-editing applications. Target DNA, defined by a pair of double strand breaks (DSBs), is removed during non-homologous end-joining (NHEJ). However, the low efficiency of CRISPR-del results in laborious experiments and false negative results. Using an endogenous reporter system, we demonstrate that temporary inhibition of DNA-dependent protein kinase (DNA-PK) – an early step in NHEJ - yields up to 17-fold increase in DNA deletion. This is observed across diverse cell lines, gene delivery methods, commercial inhibitors and guide RNAs, including those that otherwise display negligible activity. Importantly, the method is compatible with pooled functional screens employing lentivirally-delivered guide RNAs. Thus, delaying the kinetics of NHEJ relative to DSB formation is a simple and effective means of enhancing CRISPR-deletion.

## Introduction

CRISPR-Cas9 technology enables a variety of loss-of-function perturbations to study the functions of genomic elements in their natural context, and engineer natural and unnatural mutations (Cong et al. 2013; Mali et al. 2013; Doench 2017). One such application, CRISPR-deletion (CRISPR-del), is a means of permanently removing specific genomic fragments from 10^1^ – 10^6^ base pairs (Canver et al. 2014). This range has enabled researchers to investigate a wide variety of functional elements, including gene regulatory sequences (Canver et al. 2015; Mochizuki et al. 2018; Gasperini et al. 2019), non-coding RNAs (Han et al. 2014; Ho et al. 2015; Holdt et al. 2016; Koirala et al. 2017; Xing et al. 2017), and structural elements (Huang et al. 2018). Similarly, engineered deletions can be used to model human mutations (Lupiañez et al. 2015; Nelson et al. 2016). CRISPR-del is readily scaled to high throughput screens, via pooled lentiviral libraries of thousands of paired single guide RNAs (sgRNAs) (Vidigal and Ventura 2015; Aparicio-Prat et al. 2015). This has been used to discover long noncoding RNAs (lncRNAs) regulating cancer cell proliferation (Zhu et al. 2016; Liu et al. 2018) and to map *cis*-regulatory regions of key protein-coding genes (Gasperini et al. 2017; Diao et al. 2017).

CRISPR-del employs a pair of CRISPR-Cas9 complexes to introduce double strand breaks (DSBs) at two sites flanking the target region. Thereafter it relies on the endogenous non-homologous end joining (NHEJ) process to repair the breaks so as to eject the intervening fragment (Yang et al. 2013; Maddalo et al. 2014; Ho et al. 2015; Vidigal and Ventura 2015). The two ends of target regions are defined by a pair of user-designed sgRNAs (Pulido-Quetglas et al. 2017). Paired sgRNAs may be delivered by transfection or viral transduction (Vidigal and Ventura 2015; Aparicio-Prat et al. 2015). Pooled screens require that both sgRNAs are encoded in a single vector to ensure their simultaneous delivery, and are typically performed under conditions of low multiplicity-of-infection (MOI), where each cell carries a single lentiviral insertion (Vidigal and Ventura 2015; Zhu et al. 2016; Gasperini et al. 2017; Esposito et al. 2019; Doench 2018).

The principal drawback of CRISPR-del is the low efficiency with which targeted alleles are deleted. Studies on cultured cells typically report efficiencies in the range 0% – 50% of alleles, and often <20% (Mandal et al. 2014; Thomas et al. 2020), similar to estimates from individual clones (Canver et al. 2014; Vidigal and Ventura 2015; Aparicio-Prat et al. 2015; Ho et al. 2015; Pulido-Quetglas et al. 2017). Indeed, a recent publication reported high variation in the efficiencies of paired sgRNA targeting the same region, including many that yielded negligible deletion (Thomas et al. 2020). Transfection typically yields greater efficiency than viral transduction, possibly due to higher sgRNA levels (Mangeot et al. 2019), but is incompatible with pooled screening. Although megabase-scale deletions have been reported (Han et al. 2014; Essletzbichler et al. 2014), deletion efficiency decreases with increasing target size (Canver et al. 2014). Homozygous knockout clones may be isolated by screening hundreds of single cells, however this is slow and laborious, and resulting clones may not be representative of the general population (Stojic et al. 2018). More important than these practical costs, is the potential impact of low deletion rates on the ability to discern *bona fide* functional effects arising from a given mutation (Thomas et al. 2020). Non-performing sgRNA pairs are a particular problem for pooled CRISPR-del screens, where they reduce statistical power and lead to false negative results. In the screen reported by Zhu at al,.less than half of pgRNAs yielded a detectable phenotype, strongly suggesting they do not efficiently delete their targets (Fig. 1A) (Zhu et al. 2016). Similar results are reported by Canver et al, where 65% of pgRNAs yielded deletion efficiency <20% (Canver et al. 2014). There is no observable correlation between phenotype and average predicted score of the two sgRNAs (Fig. 1B) (Zhu et al. 2016). This suggest that deletion efficiency of a given pgRNA does not simply depend on the aggregate quality of its two individual sgRNAs. The outcome is that researchers are forced to increase the coverage of deletion constructs per target, resulting in lower candidate numbers and increased costs (Doench et al. 2016; Sanson et al. 2018). Consequently, any method to improve CRISPR-del efficiency would streamline experiments and enable the discovery of presently-overlooked functional elements.

**Figure 1.**
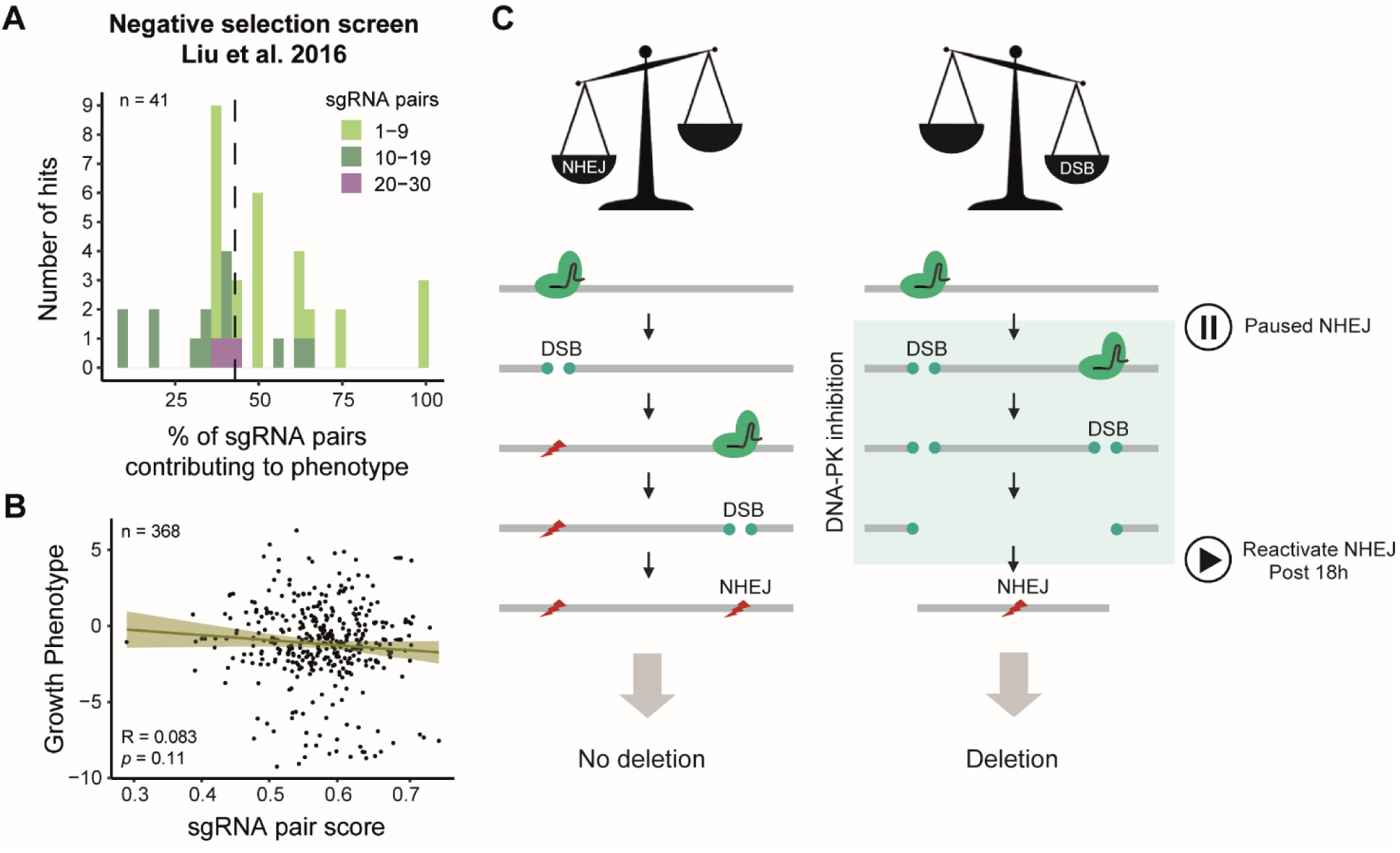
Analysis of sgRNA pairs efficiencies from public data and CRISPR-deletion model. (A, B), Re-analysis of the hits obtained CRISPR-del negative selection screen in Huh.7 cells from Zhu et al. 2016 (excluding hits targeting ORFs). (A) The upper histogram shows the percentage of sgRNA pairs contributing to the phenotype of each hit (x axis) versus the number of hits included in each bin (y axis). The dashed line represents the median (42.86%). Hits have been divided in three groups by the total number of sgRNAs designed (light green, dark green and purple). (B) The scatter plot represents the mean of the individual sgRNAs scores within a pair (calculated with the Rule Set II algorithm from Doench et al. 2016) versus the Log2 Fold Change (Growth Phenotype) obtained by Zhu et al. 2016. Pearson correlation (R) and *p*-value (*p*) are shown. (C) Model for CRISPR-del and its improvement by inhibition of DNA-PK.

For other applications of CRISPR, most notably precise genome editing using homologous recombination (HR), substantial gains have been made editing efficiency (Yeh et al. 2019). Here, editing events are rare, and HR is the rate-limiting-step (Mao et al. 2008; Miyaoka et al. 2016). The two principal strategies to boost efficiency are: (1) direct stimulation of homology directed repair (HDR) (Riesenberg and Maricic 2018; Yeh et al. 2019; Song et al. 2016; Lin et al. 2014); (2) suppression of the competing NHEJ pathway at early stages through inhibition of Ku70/80 complex (Fattah et al. 2008; Riesenberg and Maricic 2018; Yeh et al. 2019) or DNA-dependent protein kinase (DNA-PK) (Robert et al. 2015; Riesenberg and Maricic 2018; Riesenberg et al. 2019; Yeh et al. 2019), or at late phases, via Ligase IV (LigIV) inhibition (Chu et al. 2015; Maruyama et al. 2015; Riesenberg and Maricic 2018; Yeh et al. 2019). To date, however, there are no reported methods for pharmacological enhancement of CRISPR-del.

Towards this aim, we consider the events necessary for successful deletion (Fig. 1C). In the presence of two DSBs, NHEJ gives rise to successful deletion. For this to occur, the DSBs must occur on a timescale shorter than that required for NHEJ. Otherwise, the first DSB is repaired by NHEJ *before* the second can occur, and deletion will not take place. Furthermore, there is a high probability that the target protospacer or protospacer adjacent motif (PAM) is mutated during NHEJ, rendering it inaccessible to the sgRNA and precluding any subsequent deletion (Canver et al. 2014).

A prediction of this model, is that successful deletion can be promoted by extending the time over which DSBs persist without being repaired, and hence increasing the likelihood that both DSBs co-occur. In other words, we hypothesise that CRISPR-del may be improved by pharmacologically slowing the rate of NHEJ during the period while DSBs are taking place. Here, we show that inhibition of DNA-PK, an early step in NHEJ, indeed improves CRISPR-del efficiency, regardless of cell type, target region, sgRNA or inhibitory molecule, and represents a practical strategy for a variety of applications including pooled library screening.

## Results

### A quantitative endogenous reporter for CRISPR-del

To identify factors capable of improving CRISPR-del efficiency, we designed a gene-based reporter system: CRISPR Deletion Endogenous Reporter (CiDER). Such a system should be quantitative, sensitive, practical and able to closely model the CRISPR-del process by targeting endogenous genes rather than plasmids. We focussed on genes encoding cell-surface proteins, as they can be rapidly and sensitively detected by flow cytometry (Bausch-Fluck et al. 2015). A number of candidates were considered with criteria of (1) non-essentiality for cell viability and proliferation (Luo et al. 2008; Meyers et al. 2017; Tsherniak et al. 2017) (https://depmap.org), (2) high expression in human cell lines (Thul et al. 2017) (http://www.proteinatlas.org), (3) lack of overlap with other genomic elements that could lead to false positive detection, and (4) availability of flow-cytometry grade antibody. Consequently, we selected *PLXND1* encoding the Plexin-D1 protein (Supplementary Fig. 1), which presents three genomic copies in HeLa cells and two in HCT116 cells according to the Cancer Cell Line Enciclopedia (CCLE) (Barretina et al. 2012) (https://portals.broadinstitute.org/ccle/data).

We conceived an experimental setup where only successful CRISPR-del leads to loss of *PLXND1* expression, but unsuccessful events do not. In this scheme, the gene’s first exon is targeted for deletion by a series of sgRNA pairs recognising the non-protein coding regions upstream (promoter) and downstream (first intron) (Fig. 2A). Successful deletions of the first exon are expected to silence protein expression, but indels from individual sgRNAs do not affect the protein sequence directly and should not lead to silencing. Finally, we also designed sgRNAs that directly target the open reading frame (ORF), since these are expected to yield maximal protein silencing (designated positive control, *P+*).

**Figure 2.**
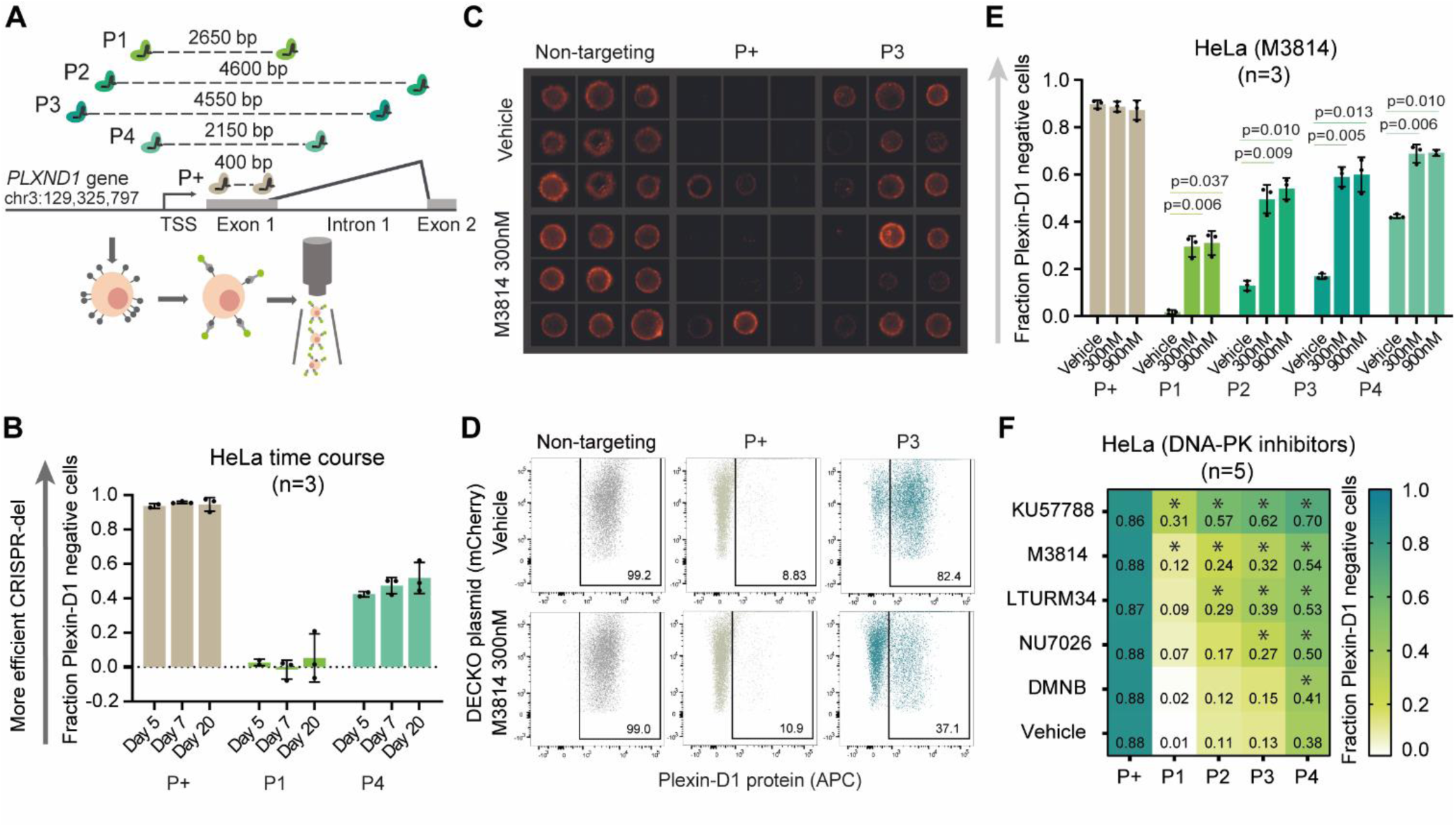
CiDER reporter system identifies DNA-PK inhibition as a means to increase CRISPR-del efficiency. (A) The CiDER endogenous reporter relies on a series of sgRNA pairs targeting exon 1 of *PLXND1* locus, whose protein product is read out by flow cytometry. (B) CRISPR-del efficiency time course in HeLa (mean, standard deviation). (C) Representative images of Plexin-Dl (APC) staining in HeLa. (D) Representative raw flow cytometry plots of CiDER in HeLa upon DNA-PK inhibition. Plexin-Dl positive cells are gated and numbers correspond to percentage of cells. (E) CRISPR-del efficiency of CiDER in HeLa upon DNA-PK inhibition (mean, standard deviation, 2-tailed paired *t*-test). (F) CRISPR-del efficiency of CiDER in HeLa upon DNA-PK inhibition with different small molecules (mean and 2-tailed paired *t*-test).

We used flow cytometry to evaluate Plexin-D1 protein levels (Fig. 2B). Positive control sgRNAs (*P+*) yielded approximately 90% knockout efficiency. We observed wide variability in the deletion efficiency of sgRNA pairs, from Pair1 (*P1*) displaying minimal efficacy, to the most efficient *P4* yielding ∼40% deletion. Therefore these paired sgRNAs achieve deletion efficiencies that are comparable to previous studies (Canver et al. 2014; Pulido-Quetglas et al. 2017). Measured deletion rates were consistent across biological replicates (Fig. 2B). The observed loss of Plexin-D1 was not due to large indels or disruption of gene regulatory elements at individual sgRNA target sites (Kosicki et al. 2018), since control experiments with single sgRNAs showed no loss of Plexin-D1 (Supplementary Fig. 2). In CiDER we have a reproducible and practical reporter of CRISPR-del at a range of efficiencies.

### Temporary inhibition of DNA-PK during DSB formation increases CRISPR-del efficiency

We hypothesized that temporarily inhibiting NHEJ during DSB formation would favour CRISPR-del, by increasing the chance that both DSBs will co-occur (Fig. 1C). We tested DNA-PK, a DNA end-binding factor at the first step of NHEJ pathway, for which a number of small-molecule inhibitors are available (Harnor et al. 2017). We began by treating HeLa cells with the inhibitor M3814 (IC_50_=3nM) (Fuchss et al. 2014; Zenke et al. 2016; Riesenberg et al. 2019) at two concentrations (300 nM and 900 nM). Importantly, cells constitutively expressing Cas9 were treated for an 18 hour time window, 4 hours after sgRNA expression plasmid delivery by transfection. Thus, DNA-PK was inhibited immediately before sgRNA expression. This resulted in improved deletion rates for all four sgRNA pairs, including a 17-fold increase for *P1*, which otherwise displays negligible deletion under normal conditions (Fig. 2C,E).

We next asked whether other inhibitors of DNA-PK yield a similar effect. We treated cells with four other commercially-available molecules at a concentration of 10uM: KU57788 (IC_50_=14nM), NU7026 (IC_50_=230nM), LTURM34 (IC_50_=34nM) and DMNB (IC_50_=15uM) (Fig. 2F). Each one yielded increases in CRISPR-del efficiency to varying degrees, correlating with published differences on the inhibition potency (Mohiuddin and Kang 2019). As expected based on previous literature, KU57788 gave the strongest effect (Mohiuddin and Kang 2019) and DMNB gave the weakest effect, likely due to its high IC_50_.

We were curious whether improved deletion depends on inhibition specifically of DNA-PK, or more generally on NHEJ. To answer this, we used SCR7 pyrazine to inhibit another step in NHEJ, the final ligation by Ligase IV (LigIV). In contrast to DNA-PK, this treatment did not improve deletion efficiency (Fig. 3A). At this late stage, the NHEJ machinery (DNA-end binding and processing complex) is already maintaining together the free DNA ends. When LigIV activity is restored, it may be more likely that each single DSB is repaired independently, introducing small indels rather than favouring genomic deletion. Thus, CRISPR-del efficiency improvements depend specifically on inhibition of DNA-PK activity. Altogether, we have shown that pharmacological inhibition of NHEJ at the DNA-PK step yields enhanced deletion of *PLXND1* reporter in HeLa cells.

**Figure 3.**
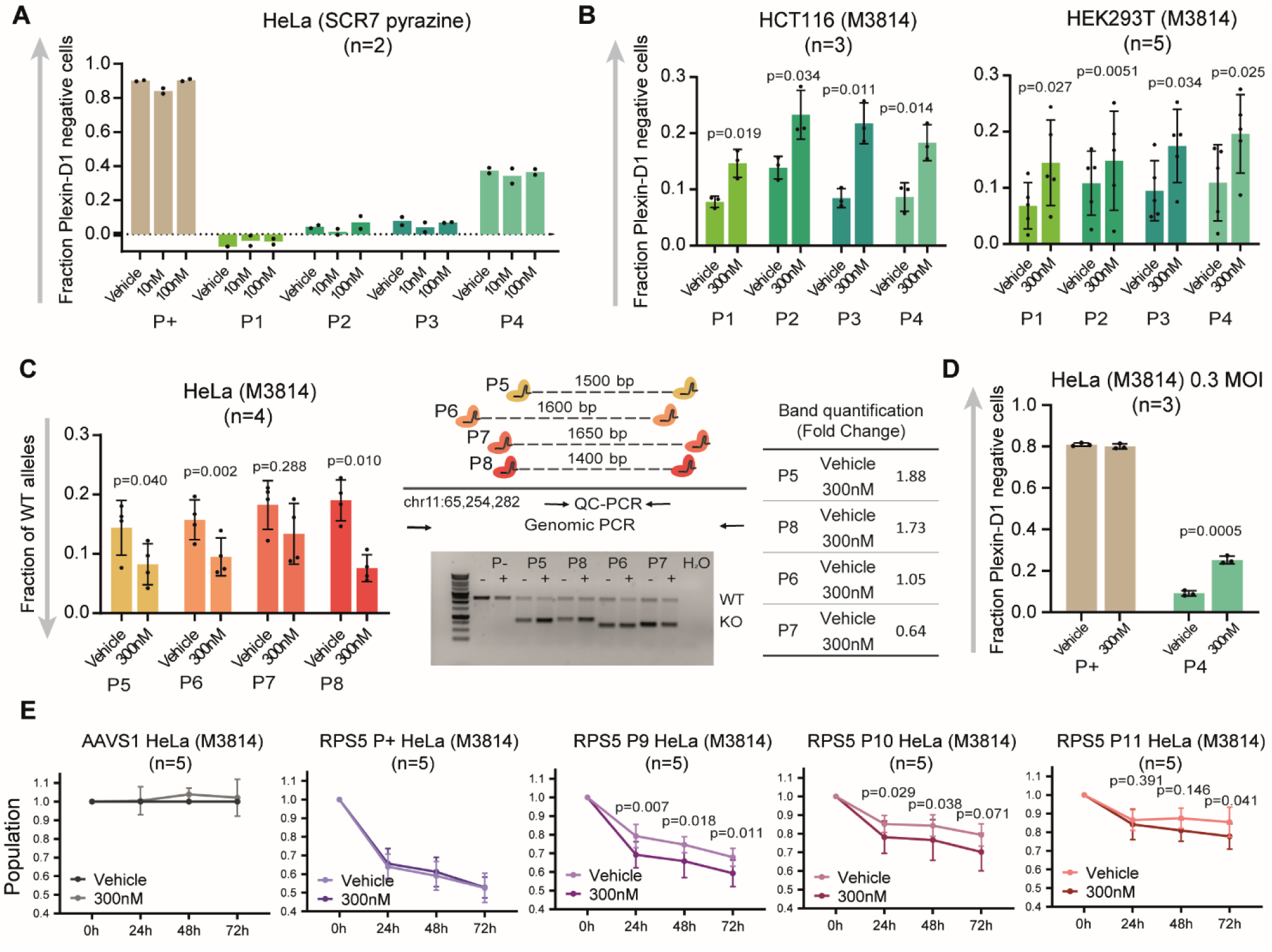
Universality of DNA-PK inhibition. (A) CRISPR-del efficiency of CiDER in HeLa upon LigIV inhibition (mean). (B) CRISPR-del efficiency of CiDER in HCT116 and HEK293T cell lines upon DNA-PK inhibition (mean, standard deviation, 2-tailed paired *t-*test). (C) CRISPR-del efficiency in chr11-locus in HeLa upon DNA-PK inhibition. The bar plots show the fraction of WT allele quantified by qPCR (mean, standard deviation, 2-tailed paired *t-*test). Shown a scheme of the sgRNA pairs and PCR primers design for this locus. Also shown a representative agarose gel from the genomic PCR of the region and band quantification of the KO allele. (D) CRISPR-del efficiency of CiDER in HeLa upon low MOI lentiviral infection and DNA-PK inhibition (mean, standard deviation, 2-tailed paired *t*-test). (E) Functional validation of DNA-PK inhibition. Viability assays after RPS5 TSS deletion upon DNA-PK inhibition (mean, standard deviation, 2-tailed paired *t*-test).

### Generality of deletion enhancement by DNA-PK inhibition

We next assessed whether this DNA-PK-inhibition is more generally effective across cell lines, genomic targets and sgRNA delivery modalities.

We began by replicating CiDER experiments in two widely-used cell lines, HCT116 and HEK293T (Thomas et al. 2020; Li and Richard 2016; Hart et al. 2015; Liu et al. 2017) (Fig. 3B). Both have baseline CRISPR-del efficiency below HeLa, possibly due to weaker NHEJ activity (Miyaoka et al. 2016). Nevertheless, DNA-PK inhibition enhanced deletion in both cell backgrounds.

All experiments so far involved a single target locus, assayed by flow cytometry. We next assessed whether these effects hold for other loci and readouts. We previously used a quantitative PCR method (quantitative CRISPR PCR, QC-PCR) to measure rates of deletion at the *MALAT1* enhancer region (Pulido-Quetglas et al. 2017). For three out of four sgRNA pairs, we observed a significant enhancement of deletion with M3814 treatment of HeLa (Fig. 3C, note the inverted scale used for QC-PCR). Similar but weaker results were also observed for HCT116 (two out of four pairs) and HEK293T cells (one out of four pairs) (Supplementary Fig. 3).

Together, although differences in performance are observed between cell types, these findings support the general applicability of DNA-PK inhibition.

### DNA-PK inhibition in the context of high-throughput pooled screens

CRISPR-del perturbations can be employed in the context of pooled functional screens, where libraries of paired sgRNAs are delivered by lentivirus at low MOI, and the effect on phenotypes such as proliferation are recorded. We asked whether DNA-PK inhibition is also practical under these conditions, by targeting the *PLXND1* reporter with sgRNAs delivered by low-MOI lentivirus. In initial experiments, M3814 was added to cell media prior to lentiviral transduction, but no improvement in deletion efficiency was observed (data not shown). This is explained by the fact that lentiviruses require NHEJ for genomic integration (Li et al. 2001; Rene et al. 2004). Therefore, we modified our protocol so as to leave sufficient time for viral integration before NHEJ inhibition (24 h was optimal, Supplementary Fig. 4), and observed a 2.7-fold increase in CRISPR-del efficiency (Fig. 3D).

Pooled CRISPR screens employ phenotypic readouts, often in the form of cell proliferation (Esposito et al. 2019; Doench 2017). To test whether improved CRISPR-del translates into stronger phenotypes, we developed a reporter assay capable of quantifying the phenotypic effect of CRISPR-del in terms of cell death. Analagous to *PLXND1* (Fig. 2A), we designed three pairs of sgRNAs targeting the first exon of the essential gene, *RPS5* (coding for the 40S ribosomal protein S5, P46782, Uniprot): *RPS5-P+, P9, P10, P11*. As expected, sgRNAs targeting the *AAVS1* locus had no effect, while sgRNAs targeting the RPS5 ORF (*RPS5-P+*) resulted in ∼47% mortality after 72 h (Fig. 3E). Neither was affected by M3814, indicating no detectable toxicity at this working concentration, as also shown by Riesenberg et al (Riesenberg et al. 2019). In contrast, three pairs of sgRNAs targeting the first exon of *RPS5* (*P9, P10, P11*) resulted in a substantial mortality (32%, 21% and 15%, respectively), which was significantly enhanced by addition of M3814 (41%, 30% and 22%, respectively).

In conclusion, DNA-PK inhibition enhances CRISPR-del when sgRNAs are delivered lentivirally at low MOI, and results in increased downstream phenotypic effects, supporting its utility in the context of high-throughput pooled screens.

## Discussion

The intrinsic DNA damage response underpins CRISPR-Cas9 genome editing and may be manipulated to favour desired editing outcomes. In the case of precise genome editing, which is based on the HDR pathway, efficiency has been substantially improved through pharmacological promotion of HDR and inhibition of the competing NHEJ pathway (Yeh et al. 2019). No such solutions have been developed for CRISPR-del, despite its being one of the most common CRISPR-Cas9 modalities, with diverse scientific and technological applications (Mochizuki et al. 2018; Holdt et al. 2016; Gasperini et al. 2019; Han et al. 2014; Ho et al. 2015; Canver et al. 2015; Koirala et al. 2017; Xing et al. 2017; Huang et al. 2018; Lupiañez et al. 2015; Nelson et al. 2016).

We hypothesised that successful CRISPR-del requires paired DSBs to co-occur *before* NHEJ has time to act, and thus may be enhanced by pharmacological inhibition of DNA-PK. This is initially counter-intuitive, as DNA-PK is a necessary step in the NHEJ pathway upon which CRISPR-del relies, and its inhibition is widely used to promote HDR(Yeh et al. 2019; Riesenberg and Maricic 2018; Riesenberg et al. 2019; Robert et al. 2015). However, rather than permanently blocking NHEJ, our protocol slows the kinetics of NHEJ for a defined period while DSBs are taking place. This produces a significant enhancement of DNA deletion efficiency, increasing protein knockout rates and resulting in stronger functional effects.

DNA-PK inhibition represents a practical option for a variety of CRISPR-del applications, from basic research to gene therapy. DNA-PK inhibitors are cheap and widely-available. Deletion efficiency improved regardless of the inhibitor molecule, target region, sgRNA sequence, cell background and delivery method. Particularly striking was the observation that some sgRNA pairs that are ineffective under normal conditions, achieved respectable rates of deletion using DNA-PK inhibition. This suggests that the failure of many sgRNA pairs to efficiently delete DNA may arise not from their inability to promote DSBs, but rather as a result of poor kinetic properties (for example, a mismatch in kinetics between the two individual sgRNAs). Finally, this method (with minor modifications) is compatible with low-MOI lentiviral delivery and leads to improvements in observed cell phenotypes. These conditions are employed in pooled screens to probe the functions of non-protein coding genomic elements (Zhu et al. 2016; Gasperini et al. 2017; Diao et al. 2017; Liu et al. 2018), meaning that DNA-PK inhibition may be used in future to improve the sensitivity of CRISPR-deletion screens by boosting the number of active sgRNA pairs, and their efficiency. This study focussed on genetic delivery methods and transformed cell types that are commonly used in high-throughput screens. Nevertheless, it will be important to assess in future the performance of DNA-PK inhibition with Cas9 ribonucleoprotein (RNP) delivery methods and in primary and non-transformed cellular backgrounds.

## Materials and methods

### Cell culture

HeLa, HCT116 and HEK293T were cultured on Dulbecco’s Modified Eagles Medium (DMEM) (Sigma-Aldrich, D5671) supplemented with 10% Fetal Bovine Serum (FBS) (ThermoFisher Scientific, 10500064), 1% L-Glutamine (ThermoFisher Scientific, 25030024), 1% Penicillin-Streptomycin (ThermoFisher Scientific, 15140122). Cells were grown at 37°C and 5% CO_2_ and passaged every two days at 1:5 dilution.

### Generation of Cas9 stable cell lines

HeLa cells were infected with lentivirus carrying the Cas9-BFP (blue fluorescent protein) vector (Addgene 52962). HCT116 and HEK293T were transfected with the same vector using Lipofectamine 2000 (ThermoFisher Scientific, 11668019). All cell types were selected with blasticidin (4ug/ml) for at least five days and selected for BFP-positive cells twice by fluorescence activated cell sorting.

### sgRNA pair design and cloning

sgRNA pairs were designed using CRISPETa (http://crispeta.crg.eu/) and cloned into the pDECKO backbone as described previously (Pulido-Quetglas et al. 2017). Off-target filters did not allow less than 3 mismatches for each sgRNA sequence. No positive or negative masks were applied in the search. Minimum individual score was set at 0.2 and minimum paired score at 0.4. The sgRNA pairs were then manually selected from the output list. All sgRNA sequences may be found in Supplementary Figure 5.

### Inhibitors

All molecules used in this study are commercially available: M3814 (MedChemExpress, HY-101570), KU57788 (MedChemExpress, HY-11006), NU7026 (MedChemExpress, HY-15719), LTURM34 (MedChemExpress, HY-101667), DMNB (ToChris, 2088) and SCR7 Pyrazine (Sigma-Aldrich, SML1546). 10mM stocks (and 5mM for NU7026, due to solubility limitations) were prepared by resuspension in dimethylsulfoxide (DMSO) (Sigma-Aldrich, D4540).

### Transfection and lentiviral transduction

For transfection experiments, 70% confluent 12-well plates were transfected using Lipofectamine 2000 (ThermoFisher Scientific, 11668019) with 1250 ng of pDECKO plasmid following provider’s guidelines. After 6 hours, transfection media was replaced for fresh complete DMEM (10% FBS, 1% L-Glutamine and 1% Penicillin-Streptomycin) and the corresponding small molecule was added to media for 18 hours. The treatment was finished by replacing the media with complete DMEM. After one day cells were selected with puromycin (2ug/ml).

For lentiviral infection experiments, cells were spin-infected at a 0.3 multiplicity of infection in the presence of DMEM (10% FBS, 1% L-Glutamine) and hexadimethrine bromide (8ug/ml) (Sigma-Aldrich, 107689) at 2000 rpm, 37°C during 1.5 hours. After 0, 5, 10, 24, 48, 72 hours, infection media was replaced for fresh complete DMEM (10% FBS, 1% L-Glutamine and 1% Penicillin-Streptomycin) and the corresponding small molecule was added to media for 18 hours. The treatment was finished by replacing the media with complete DMEM and puromycin (2ug/ml) to start the selection.

### Flow cytometry

After five days of puromycin selection, cells were trypsinized, resuspended in PBS and incubated for 30 minutes at room temperature (RT) with the human α-PlexinD1 mouse monoclonal antibody (1:150 dilution) (R&D systems, MAB4160). Cells were washed twice with PBS and incubated for 30 minutes at RT with an α-Mouse IgG secondary goat antibody conjugated to the APC fluorochrome (1:200 dilution) (eBioscience, 17-4010-82). Cells were washed and resuspended in PBS, processed with the LSRII SORP flow cytometer and analysed with FlowJo_v10 software. A total of 10,000 cells per sample are sorted. Cell population is selected in the SSC-A/FSC-A plot. Single cells are gated in the FSC-H/FSC-A plot. Finally, the APC positive population is set in the mCherry/APC plot in the control sample and expanded to all the other samples without modification. The fraction of Plexin-D1 negative singlet cells is calculated by gating Plexin-D1 positive singlet cells, normalizing to a non-targeting control and subtracting the value to 1 (negative cells = 1 – positive cells). An example of the gating strategy may be found in Supplementary Figure 6.

Single cell imaging was performed using ImageStream (Luminex) and analysed with IDEAS software.

### Genomic PCRs

After 5 days of puromycin selection, cells were collected and genomic DNA (qDNA) was extracted using GeneJET Genomic DNA Purification Kit (ThermoFisher Scientific, K0722). Genomic PCR was performed using GoTaq® G2 DNA Polymerase (Promega, M7841) from 10ng gDNA (Forward: *5’ CCTGCTATGAACTGACCCATG 3’*, Reverse: *5’ CCTGAACAGTCAGTCCATGCT 3’*)

### Genomic quantitative PCRs

After 5 days of puromycin selection, cells were collected and genomic DNA (qDNA) was extracted using GeneJET Genomic DNA Purification Kit (ThermoFisher Scientific, K0722). Quantitative real time PCR (qPCR) from 10ng of gDNA was performed using GoTaq qPCR Master Mix (Promega, A6001) on a TaqMan Viia 7 Real-Time PCR System. (Target sequence - Forward: *5’ GCTGGGGAATCCACAGAGAC 3’*, Reverse: *5’ CATCTCAGCCCTTGTTATCCTG 3’*) and (LDHA - Forward: *5’ TGGGCAGTAGAAAGTGCAG 3’*, Reverse: *5’ TACCAGCTCCCACTCACAG 3’*). Target sequence primers were normalized to primers targeting the distal, non-targeted gene LDHA. Data were normalised using the ΔΔCt method (Schmittgen and Livak 2008).

### Cell viability assay

CellTiter-Glo® 2.0 Cell Viability Assay (Promega, G9241) was performed upon puromycin selection (2 days post transfection). 3000 cells/well were seeded in 96-well white polystyrene plates (Corning®, Sigma-Aldrich CLS3610-48EA) and cell viability was measured in technical duplicates during 4 consecutive days (0h, 24h, 48h, 72h) according to the manufacturer’s protocol. Luminescence was measured using a Tecan Reader Infinite 200.

## Acknowledgements

We gratefully acknowledge administrative support from Ana Radovanovic and Silvia Roesselet (DBMR, University of Bern). We thank Bill Keyes (IGBMC) and Norbert Polacek (DCB, University of Bern) for insightful feedback and discussions. We also thank Stefan Müller (DBMR, University of Bern) for his expertise with ImageStream and the other members of the FACS lab from the University of Bern for their advice. We also acknowledge Taisia Polidori and Paulina Schaerer (DBMR, University of Bern) for the experimental support, Álvaro Andrades (Universidad de Granada) for the CCLE data and the rest of the members of Johnson’s lab for their valuable input. Andrea Maddalena (Department of Physiology, University of Bern) provided valuable advice regarding lentiviral transduction inhibition.

## Author contributions

N.B. and R.J. conceived and designed the experiments. N.B. performed all the experiments and analysis of public data. M.M. and Y.Z. suggested DNA-PK inhibition to modulate NHEJ. C.P. contributed on the design of the sgRNA pairs and analysis of public data. R.E. provided the solution to circumvent lentiviral infection problems. N.B. and R.J. wrote the whole manuscript with feedback from M.M. and Y.Z. Finally, R.J. directed the research.

## Data availability

The authors declare that all the data supporting the findings of this study are available within the paper and its supplementary information files.

## Ethics approval and consent to participate

Not applicable.

