## Supplementary Figures for "Enhancing CRISPR deletion via pharmacological delay of DNA-PK"

a

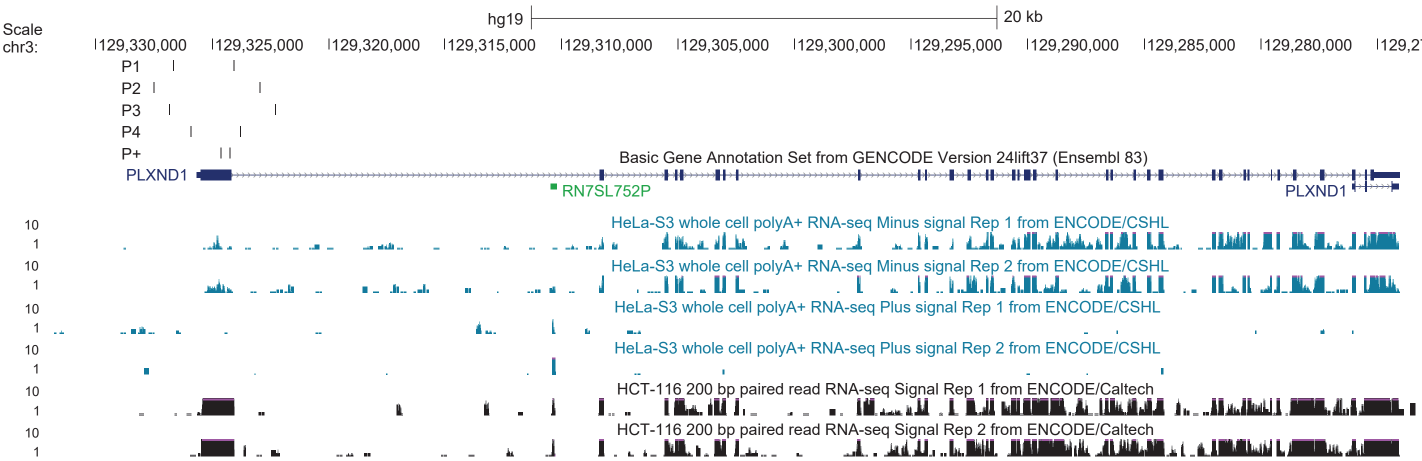


**Supplementary Fig. 1| *PLXND1* genomic locus. a,** Image from the UCSC Genome Browser (GRCh37/hg19). *PLXND1* gene is shown in dark blue (exons are represented as thick boxes and introns as lines). sgRNA pairs are depicted (*P1*, *P2*, *P3*, *P4*, *P+*) around exon 1. Whole-cell RNA expression in HeLa (ENCODE/Cold Spring Harbor Lab; plus and minus strand) and HCT116 (ENCODE/Caltech).

**a**

HeLa (M3814)
(n=3)

Fraction of Plexin-D1 negative cells

| | | Average | Standard deviation |
|---|---|---|---|
| P+ | Vehicle | 0.909 | 0.013 |
| | 300nM | 0.896 | 0.017 |
| P1.1 | Vehicle | 0.013 | 0.015 |
| | 300nM | 0.000 | 0.008 |
| P1.2 | Vehicle | 0.009 | 0.005 |
| | 300nM | 0.004 | 0.007 |
| P2.1 | Vehicle | 0.004 | 0.011 |
| | 300nM | 0.002 | 0.009 |
| P2.2 | Vehicle | 0.006 | 0.009 |
| | 300nM | 0.003 | 0.009 |
| P3.1 | Vehicle | 0.006 | 0.012 |
| | 300nM | 0.002 | 0.007 |
| P3.2 | Vehicle | 0.001 | 0.009 |
| | 300nM | 0.000 | 0.008 |
| P4.1 | Vehicle | 0.016 | 0.015 |
| | 300nM | 0.012 | 0.012 |
| P4.2 | Vehicle | 0.016 | 0.014 |
| | 300nM | 0.010 | 0.008 |

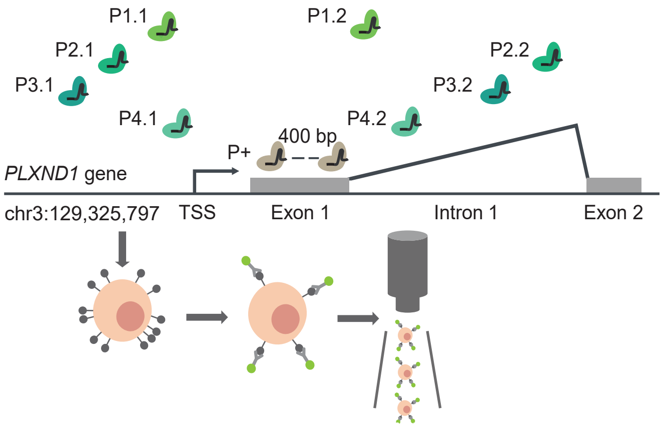


**Supplementary Fig. 2| Effect of single sgRNAs around *PLXND1* first exon. a,** CRISPR-del efficiency of each single sgRNA targeting CiDER in HeLa with and without DNA-PK inhibition (mean, standard deviation). Scheme of CiDER single sgRNAs.

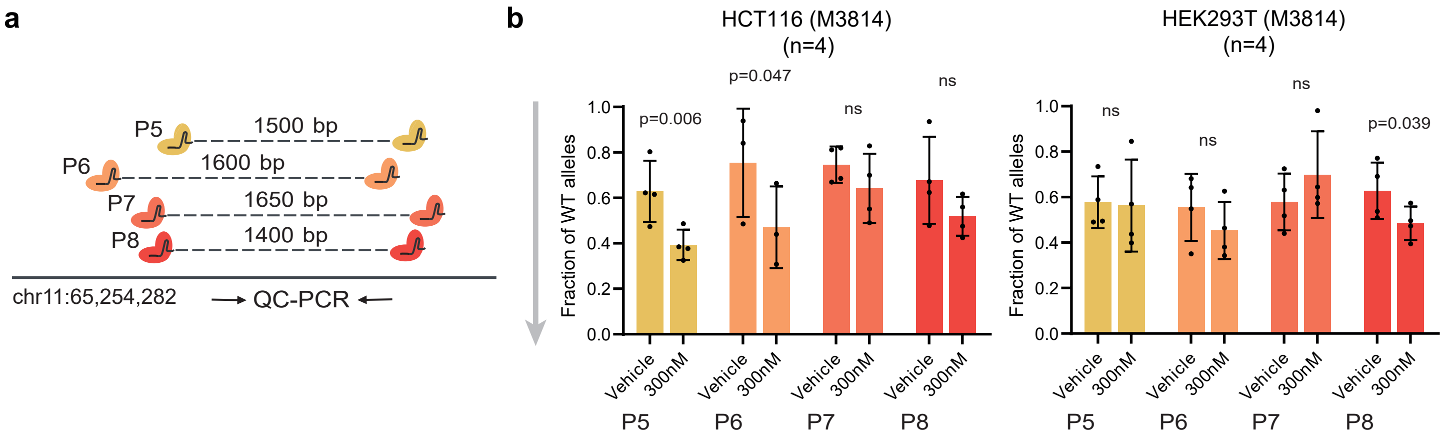


**Supplementary Fig. 3| Effect of DNA-PK inhibition in the *MALAT1* enhancer locus. a,** Scheme of the sgRNA pairs and QC-PCR primers. **b,** CRISPR-del efficiency in chr11-locus in HCT116 and HEK293T upon DNA-PK inhibition. The bar plots show the fraction of WT allele quantified by qPCR (mean, standard deviation, 2-tailed paired *t*-test).

**a**

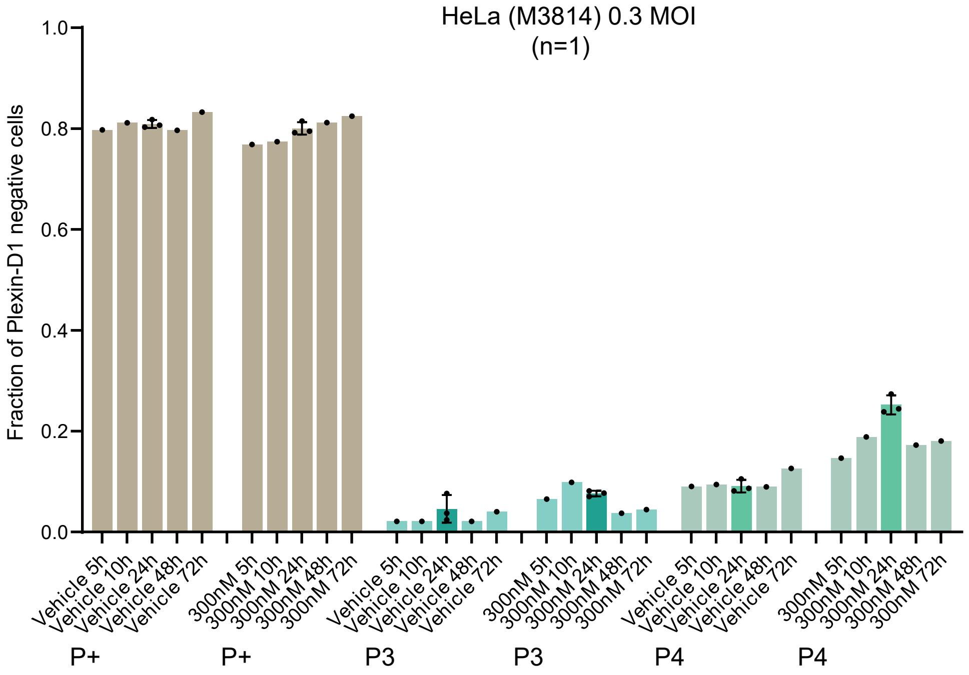


**Supplementary Fig. 4| Effect of DNA-PK inhibition in high-throughput pooled screens conditions. a,** CRISPR-del efficiency of CiDER in HeLa starting the DNA-PK inhibition in different timings post lentivirus low MOI infection (mean, standard deviation, 2-tailed paired *t*-test).

**a**

| | chromosome | start | end | sgRNA_1 | score_1 | chromosome | start | end | sgRNA_2 | score_2 | distance | paired_score |
|---|---|---|---|---|---|---|---|---|---|---|---|---|
| *PLXND1* P+ | chr3 | 129324592 | 129324615 | GCGTAGGACTGTGCACCCGG | 0.76 | chr3 | 129324215 | 129324238 | GGTGGCGGTGCTCGACAGCG | 0.74 | 400 | 1.50 |
| P1 | chr3 | 129324014 | 129324037 | GTATTCTCGCGTGCACACCT | 0.74 | chr3 | 129326648 | 129326671 | TGATCTCAAAAGCAGCGTTA | 0.22 | 2611 | 0.96 |
| P2 | chr3 | 129327478 | 129327501 | ATAGGAACAGAGATGGGTGG | 0.80 | chr3 | 129322928 | 129322951 | CATGTGTGTTGACATGACAA | 0.74 | 4527 | 1.55 |
| P3 | chr3 | 129326800 | 129326823 | TTTCCTCACAGATTCCCCCG | 0.82 | chr3 | 129322248 | 129322271 | ACAGATCAGCTTACACCAAA | 0.77 | 4529 | 1.59 |
| P4 | chr3 | 129323746 | 129323769 | TCAGAGTTACCATTGCACGT | 0.81 | chr3 | 129325881 | 129325904 | CGCGAGCGAGGCAGGCCAAG | 0.39 | 2112 | 1.21 |
| P5 | chr11 | 65254685 | 65254708 | GTTGGTCAAGTAAAGACACG | 0.84 | chr11 | 65256189 | 65256212 | AGTTCTGCCTCAGCTCAGGA | 0.68 | 1481 | 1.52 |
| P6 | chr11 | 65254522 | 65254545 | GCAGGTCACGCAGGCCCCCA | 0.44 | chr11 | 65256189 | 65256212 | AGTTCTGCCTCAGCTCAGGA | 0.68 | 1644 | 1.13 |
| P7 | chr11 | 65254522 | 65254545 | GCAGGTCACGCAGGCCCCCA | 0.44 | chr11 | 65256126 | 65256149 | GCACCAGCCCAAGGCTGCAT | 0.44 | 1581 | 0.88 |
| P8 | chr11 | 65254685 | 65254708 | GTTGGTCAAGTAAAGACACG | 0.84 | chr11 | 65256126 | 65256149 | GCACCAGCCCAAGGCTGCAT | 0.44 | 1418 | 1.27 |
| *RPS5* P+ | chr19 | 58904535 | 58904558 | GACCTGCTCACAGGCGAGGTA | 0.53 | chr19 | 58904833 | 58904856 | ACCTGGTTCACACGGCGCAG | 0.58 | 321 | 1.12 |
| P9 | chr19 | 58897261 | 58897284 | GCAGAACTGGGACTTTCCAG | 0.85 | chr19 | 58900417 | 58900440 | GTACGGAGTAGGAACAAAGT | 0.59 | 3133 | 1.44 |
| P10 | chr19 | 58897115 | 58897138 | CCTCCAACTCGGGATCCGAA | 0.86 | chr19 | 58899083 | 58899106 | TTATATGACATCAAGGACCA | 0.76 | 1945 | 1.62 |
| P11 | chr19 | 58897115 | 58897138 | CCTCCAACTCGGGATCCGAA | 0.86 | chr19 | 58900434 | 58900457 | AGGGTATGTGTAAAACCGTA | 0.59 | 3296 | 1.45 |

**Supplementary Fig. 5| Details of all the sgRNA pairs. a,** Table indicating the genomic location and sequence of each sgRNA pair and their scores (individual and paired) from CRISPETa. Distance between each sgRNA within a pair is also shown.

a

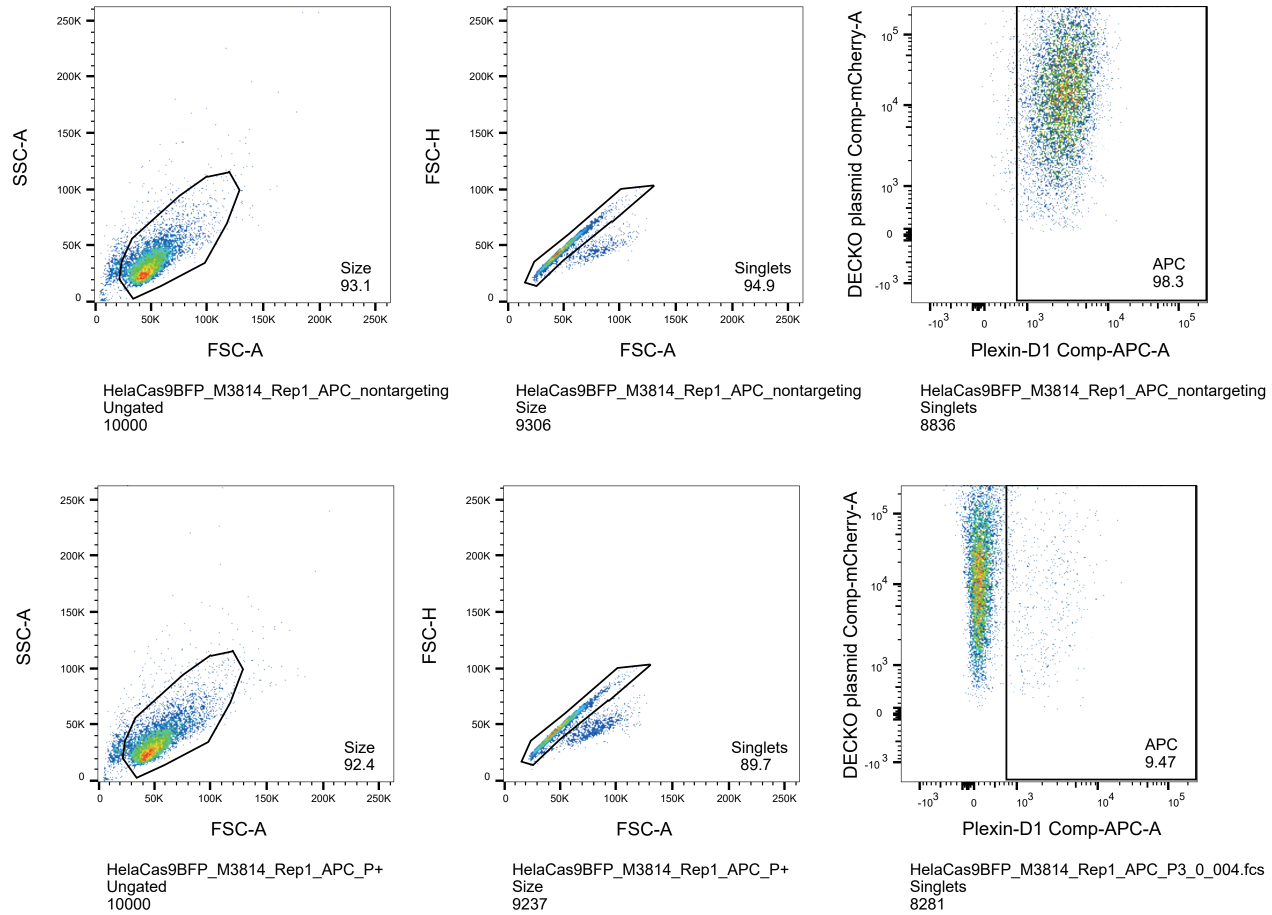


**Supplementary Fig. 6| Representative example of the gating strategy. a,** First column shows the gating for initial cell population, second column the gating for single cells and third column the gating for Plexin-D1 positive cells (shown the percentage of cells in respect to the parental population). On the bottom of each plot is shown the name of the sample, the parental population and the number of cells in each population.
